# Lipid Droplets and Ferritin Heavy Chain: a Devilish Liaison in Cancer Radioresistance

**DOI:** 10.1101/2021.05.11.443559

**Authors:** L. Tirinato, M.G. Marafioti, F. Pagliari, J. Jansen, I. Aversa, R. Hanley, C. Nisticò, D. Garcia-Calderón, G. Genard, J. F. Guerreiro, F.S. Costanzo, J. Seco

## Abstract

Although much progress has been made in cancer treatment, the molecular mechanisms underlying cancer radioresistance (RR) as well as the biological characteristic of radioresistant cancer cells still need to be clarified. In this regard, we discovered that breast, bladder, lung, neuroglioma and prostate 6 Gy X-ray resistant cells were characterized by an increase of Lipid Droplet (LD) number and that the cells containing highest LDs showed the highest clonogenic potential after irradiation. Moreover, we observed that LD content was tightly connected with the iron metabolism and in particular with the presence of the ferritin heavy chain (FTH1). In fact, breast and lung cancer cells silenced for the FTH1 gene showed a reduction in the LD numbers and, by consequence, became radiosensitive. FTH1 restoration as well as iron-chelating treatment by Deferoxamine were able to restore the LD amount and RR. Overall, these results provide evidence of a novel molecular mechanism behind RR in which LDs and FTH1 are tightly connected to each other, a synergistic effect which might be worth deeply investigating in order to make cancer cells more radiosensitive and improve the efficacy of radiation treatments.

## Introduction

Since its first application in cancer treatment, radiotherapy has greatly improved from both a technical and a bio-clinical point of view, significantly increasing the treatment options and patient survival. Ionizing radiations (X-rays) work by damaging cell biomolecules, mostly DNA, which eventually induce cell death. The molecular mechanisms activated by cancer cells in response to ionizing radiation are extensively investigated and many advances have been so far made, but considerably many questions are still unanswered and much remains poorly understood. Cancer cell radioresistance (RR) makes different tumor types difficult to treat. In this regard, the presence within the tumor mass of a small cell subpopulation called Cancer Stem Cells (CSCs) or Cancer Initiating-Cells (CICs) seems to represent one of the driving forces contributing to tumor resistance and recurrence after radiotherapy treatments ^1^.

Recently, lipid metabolic reprogramming in cancer cells has become a central aspect of cancer aggressiveness ^2,3^. In particular, an increase of small lipid organelles inside cancer cells, namely lipid droplets (LDs), has been shown to correlate with a CSC phenotype in colon ^4^ ovary ^5^, breast ^6^ and glioblastoma ^7^.

Cell survival upon radiation treatment is also modulated by several tumor parameters such as hypoxia, oxidative stress, inflammation, acidic stress, and low glucose, all of which have been reported to mediate their effects through iron metabolism ^8^.

To date, altered expression and activity of many iron-related proteins in cancer cells have been reported and associated to cancer progression and metastasis ^9,10^. In fact, an uncontrolled balance of iron results in the free radical production through the Fenton reaction (Fe^2+^ + H_2_O_2_ → Fe^3+^ + *OH + OH^-^) and free radicals are considered strong contributors to tumor proliferation and aggressiveness ^11^. Among all molecules involved in iron metabolism, ferritin is responsible for the cytoplasmic iron storage and the maintenance of the redox homeostasis. Ferritin is a protein complex composed of two chains, light (FTL) and heavy (FTH), and its clinical importance has been demonstrated in many cancers through multiple roles: the contribution to tumorigenesis, the restoration of tumor-dependent vessel growth and the association with tissue invasion ^8,12^. Moreover, high levels of ferritin are often found in patients with various advanced cancers which could potentially be treated with radiotherapy ^13^, although iron homeostasis is still poorly investigated in the context of radiation oncology.

A recently published paper highlighted a very intriguing relationship between iron balance and LDs. The Authors showed that iron depletion caused ER expansion and, as a consequence, LD accumulation into the cytoplasm of breast cancer cells ^14^. These findings prompted us to investigate the LD role and the potential connections between FTH1, and indirectly iron balance, and LDs in various X-Ray-treated cancer cells with the aim at identifying possible shared features which can be targeted and manipulated to sensitize cells to the treatments.

This study demonstrates that radioresistant cancer cells of different origin (neuroglioma, lung, breast, bladder and prostate) were characterized by a higher expression of LDs. The subpopulation containing the highest amount of LDs (LD^High^) showed a higher clonogenic potential compared to the LD^Low^ counterpart. Interestingly, the number of cytoplasmic LDs was directly correlated with the amount of FTH1. In fact, FTH1 knockdown in lung H460 (H460^shFTH1^) and breast MCF7 (MCF7^shFTH1^) reduced the LD amount and increased the sensitivity to ionizing radiation. FTH1 restoration as well as the treatment with an iron chelating agent in MCF7^shFTH1^ and H460^shFTH1^ restored the LD amount and increased their resistance to radiation treatment.

Altogether, these data provide evidence of a new pivotal role for LDs in cancer RR linking their expression with iron metabolism and specifically to FTH1 expression.

## Results

### X-ray radiation treatment enhances Lipid Droplets

To verify whether LD content was affected by ionizing radiation treatment, H4 (neuroglioma), H460 (lung), MCF7 (breast), PC3 (prostate) and T24 (bladder) cancer cells were treated with 6 Gy X-ray and left in culture for 72 hours (hrs) in order to select only resistant/surviving cells. Treated and untreated cancer cells were stained with LD540 and imaged at the confocal microscope for the detection of LDs.

As shown by z-projection confocal microscopy images, surviving cancer cells were characterized by a significant increase of LDs for all the aforementioned cell lines (**Figure 1A**). Although the LD increase was a common feature observed in all cell lines, the relative LD ratio between treated and untreated cells showed little differences, with H460 exhibiting the highest amount (**Figure 1B**). LD modulation after radiation was further investigated at the gene level. Perilipin (PLIN) genes code for the proteins associated with LD surface and they are involved in their biogenesis as well as in several other roles ^15^. Differences in tissue expression have been reported for all the PLIN genes (PLIN 1-5). Accordingly, we observed that, after 6 Gy radiation treatment, PLIN1 was up-regulated in H460, MCF7, PC3 and T24; PLIN2 was down-regulated in H460; PLIN3 showed mRNA increased expression in H4 and MCF7; PLIN4 expression was incremented in MCF7 and T24; PLIN5 resulted down-regulated in MCF7 and up-regulated in PC3 and T24.

**Figure 1:**
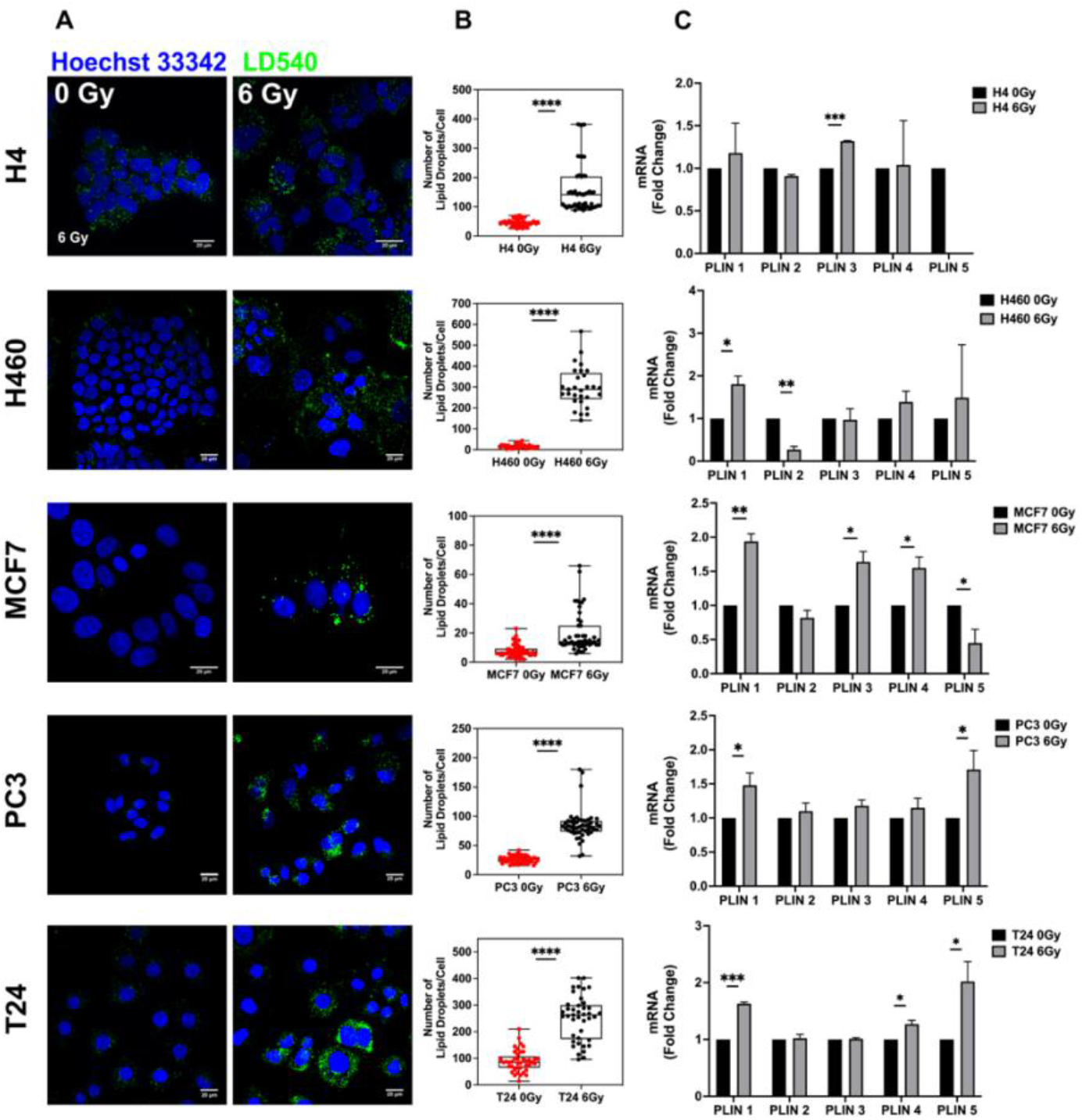
Lipid Droplet detection in neuroglioma (H4), lung (H460), breast (MCF7), prostate (PC3) and bladder (T24) 6 Gy X-ray resistant cancer cells. Cancer cells have been irradiated with 6 Gy X-ray and left in culture for 72 hrs. Afterwards, surviving and untreated cells have been stained with LD540 and imaged at the fluorescence confocal microscope. Z-projection of the z-stack acquisitions for untreated and 6 Gy treated cells are reported in column **A** (Scale bar, 20 μM). **B**) For each cell line, 50 cells have been randomly imaged, and their LD number counted by using FiJi software. **C**) qPCR analysis of the PLIN genes in the indicated cell lines. PLIN5 in the H4 6 Gy treated cells is not reported in the graph because it was not expressed. Error bars represent the means ± SD from three independent experiments. * ≤ 0.05; ** ≤ 0.01; *** ≤ 0.001 and ****≤ 0.0001.

It is well known that photon radiation acts, at the molecular level, producing reactive oxygen species (ROS) ^8^. In this regard, we found that cytoplasmatic ROS, measured by means of fluorogenic CM-H2DCFDA probe, were significantly upregulated in H4, H460, MCF7 and PC3, while no differences were detected in T24, after radiation (**Figure S1**). Moreover, H4, H460 and PC3 showed upregulated levels of SOD1, SOD2 and catalase, respectively. SOD2 mRNA was also upregulated in T24, despite the fact that general ROS levels resulted not altered 72 hrs after radiation treatment, while it was downregulated in radiation treated MCF7.

In order to deal with this ROS increase, cancer cells need to tune their ROS scavenging systems ^16^, and among all scavenging systems, LDs have been observed to contribute to the modulation of excessive oxidative stress ^17^. Furthermore, by co-staining LDs and ROS in the heterogeneous not irradiated cancer populations, we found that populations with higher LDs also exhibited higher levels of ROS (**Figure S2**). Therefore, LD content, influencing cell survival, was directly correlated with ROS production in all cell lines.

Previous works reported that ionizing radiation could selectively enrich the cancer cell population of cells with stem-like properties ^18-20^. Thus, we have analyzed the expression of some of the most common markers used to identify CSCs. In particular, we found that CD44 was upregulated in 6 Gy treated H4, H460, MCF7 and T24; CD133 mRNA increased in H4 and H460-irradiated cells; CD166 expression was upregulated in MCF7 and T24; ALDH1 was incremented in MCF7. On the contrary, PC3 RR cells did not display significant increase in the expression of such CSC markers (**Figure S3**).

### LD^high^ sub-population retains the highest clonogenic potential

LD modulation following X-ray treatment raised the question if LD accumulation was a consequence of radiation treatment or if such a feature was already present in some cells within the heterogeneous cancer populations, thus suggesting that LD content could participate in conferring a higher radiation resistance.

To better define the role played by LDs in RR cells and to address the question, H4, H460, MCF7, PC3 and T24 were stained with LD540, sorted in the 10% highest and lowest LD-containing cells (LD^High^ and LD^Low^ cells) (**Figure 2**) and, soon after, irradiated with 2, 4 and 6 Gy X-ray. The surviving fractions (SFs), calculated for all cell lines at the different doses, showed that LD^High^ cells retained the highest clonogenic potential and therefore they were the most radioresistant (**Figure 2**). These results suggest that the LD amount present in the population is linked to a stronger cell capability to survive ionizing radiations, independently of the tissue of origin.

**Figure 2:**
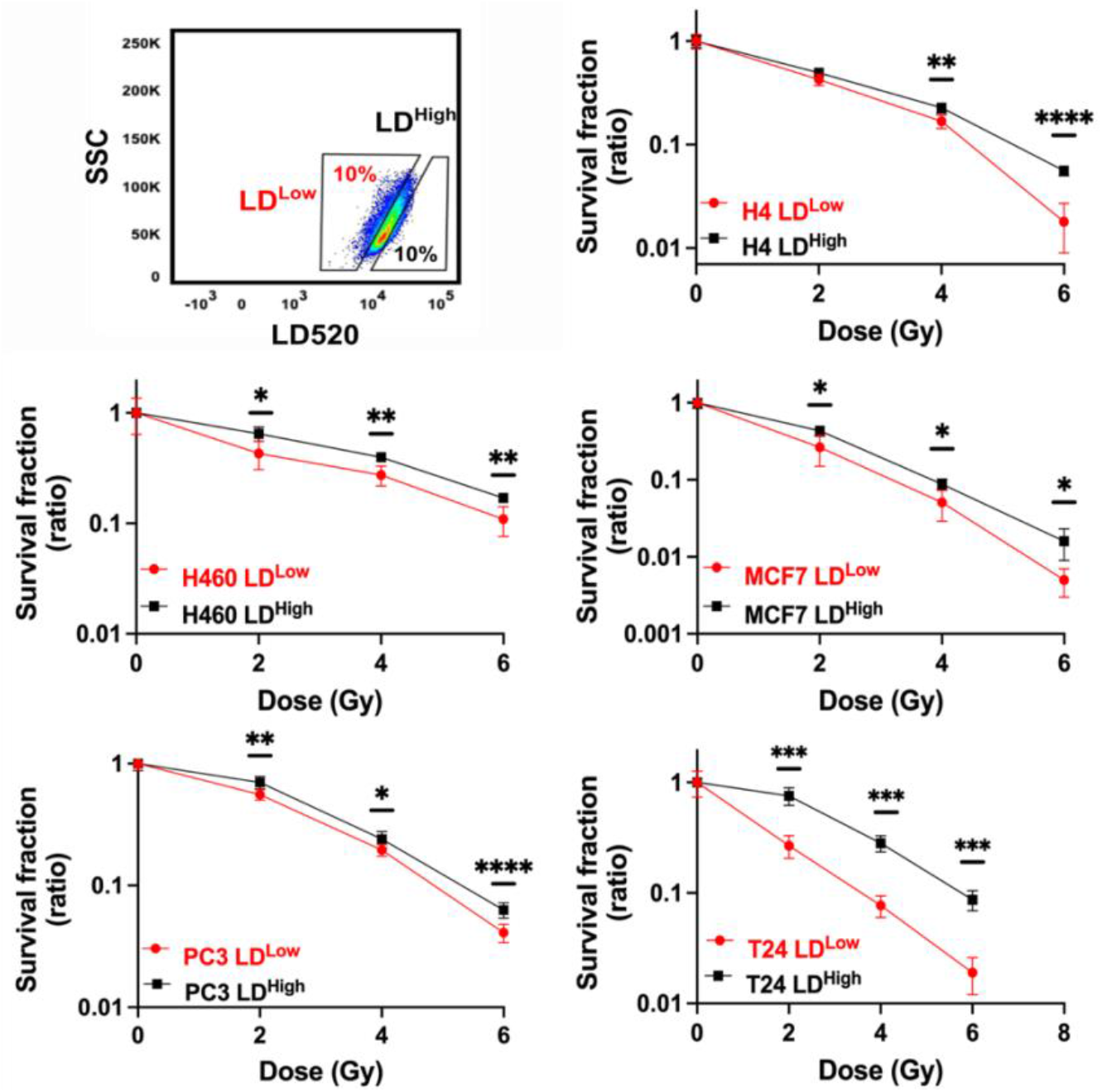
Cell survival curves for H4, H460, MCF7, PC3 and T24 cancer cell lines. All cancer cell lines were stained with LD540 and then sorted in the 10% highest and lowest LD-expressing cells (**box up-left**). For each cell line, the two LD sub-populations were irradiated at 0, 2, 4 and 6 Gy X-ray and their survival fraction calculated. Survival fractions are reported in log-linear scale. Error bar represents the means ± SD from three independent experiments. * ≤ 0.05; ** ≤ 0.01; *** ≤ 0.001 and ****≤ 0.0001.

### Ferritin Heavy Chain (FTH1) affects LD accumulation and cell radio-response

One of the main cellular ROS sources is the Fenton reaction, in which the Fe^2+^ reacts with hydrogen peroxide (H_2_O_2_) to produce Fe^3+^ and highly reactive radicals, such as the hydroxyl radical (·OH). Since the Ferritin is the main intracellular iron storage protein, we investigated the FTH1 role in radiation resistance.

We found that FTH1 protein was upregulated in all resistant cell lines after 72 hrs from 6 Gy exposure, as reported in **Figure 3A**. Moreover, H460 and MCF7, sorted on the basis of their LD content, were characterized by an increase in the mRNA level of FTH1 in the LD^high^ subpopulation compared to the LD^low^ cells (**Figure 3B**).

**Figure 3:**
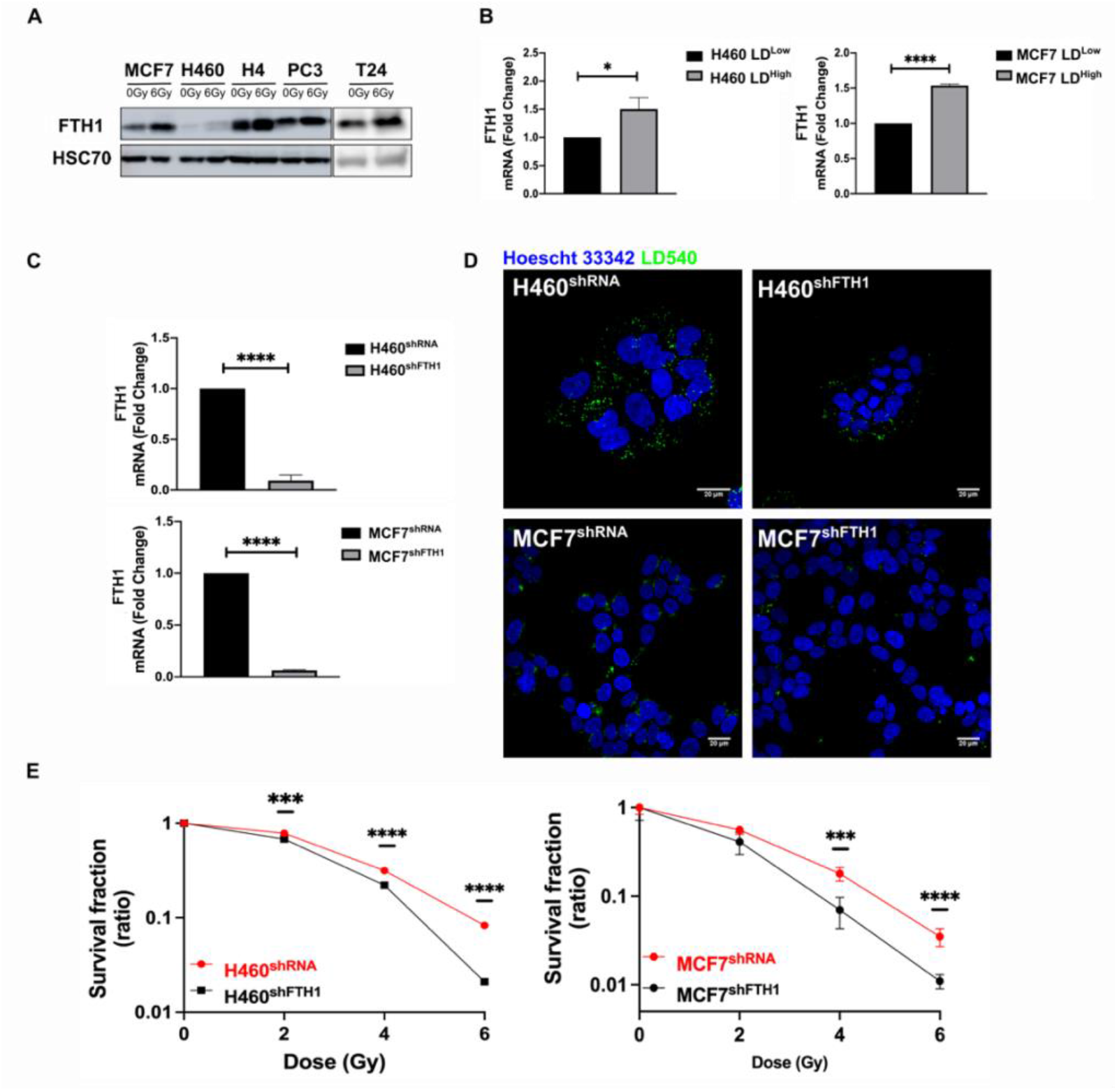
FTH1 silencing downregulates Lipid Droplets affecting cancer radioresistance. **A**) Wester blotting analysis of FTH1 expression in MCF7-, H460-, H4-, PC3-, T24-0Gy vs 6Gy X-ray treated cells. HSC70 was used as a loading control. **B**) H460 and MCF7 were sorted in the 10% Highest (H460 LD^High^ and MCF7 LD^High^) and Lowest (H460 LD^Low^ and MCF7 LD^Low^) LD-expressing cells and then FTH1 mRNA expression measured by qRT-PCR in all four sub-populations. Primer sequences are listed in the Supporting Information. **C**) H460 and MCF7 were silenced for FTH1 by lentiviral-driven shRNA strategy. PCR results showed that in H460 shFTH1 and MCF shFTH1 there was a clear FTH1 mRNA reduction compared with their relative controls. **D**) LD content was measured in H460 shFTH1 and MCF7 shFTH1 by confocal microscopy. LD540 staining revealed that the FTH1 gene silencing caused a LD decrease in both cell systems. (Scale bars 20 μM). **E**) Cellular irradiation response in H460 and MCF7 silenced for FTH1 was investigated by radiobiological clonogenic assay and compared with H460 shRNA and MCF7 shRNA respectively. Survival fraction (in log-linear scale) is reported in the panel **E**. Error bar represents the means ± SD from three independent experiments. * ≤ 0.05; ** ≤ 0.01; *** ≤ 0.001 and ****≤ 0.0001.

To better clarify this link, FTH1 was silenced in H460 and MCF7 (H460^shFTH1^ and MCF7^shFTH1)^, the efficiency of which is shown in **Figure 3C**. FTH1 silencing resulted in influencing cell ability to deal with free cytoplasmic iron, as demonstrated by the downregulation of Transferrin Receptor 1 (TfR1) mRNA and the up-regulation of Ferroportin (FPN) mRNA, all involved in proper iron homeostasis (**Figure S4 A** and **B**).

Interestingly, in FTH1 silenced H460 and MCF7, the amount of FTH1 directly correlated with the number of LDs (**Figure 3D**). In fact, H460 ^shFTH1^ and MCF^shFTH1^ cells were characterized by a significant reduction of LDs. This, in turn, led to an evident increase in the radiosensitivity, as demonstrated by the clonogenic assay results (**Figure 3E**).

Summarizing, we show that LD content was dependent on the FTH1 expression and thus linked to the free cytoplasmic iron. When the levels of the main protein responsible for iron storage decreased, LDs were also reduced and this significantly impaired cancer RR.

### Iron Imbalance as well as FTH1 reconstitution re-establish LD expression and radiation resistance

As well known, the FTH1 role is crucial for the iron storage within the cell and the maintenance of the redox homeostasis. When its expression is downregulated, the balance between the iron uptake and release is compromised. By consequence, the free cellular iron amount becomes critical for the correct cellular functions ^10,21^. Here we found that this iron imbalance assumed also a central role in the LD accumulation. Given the role played by the FTH1 deficiency on LD content and radiosensitivity, we reconstituted the FTH1 expression by full length FTH1 cDNA transfection to further verify such connection. **Figure 4A** shows that FTH1 gene restoration successfully raised FTH1 protein expression up in both MCF7^shFTH1^ + pcDNA_3_FTH1 and H460^shFTH1^ + pcDNA_3_FTH1.

**Figure 4:**
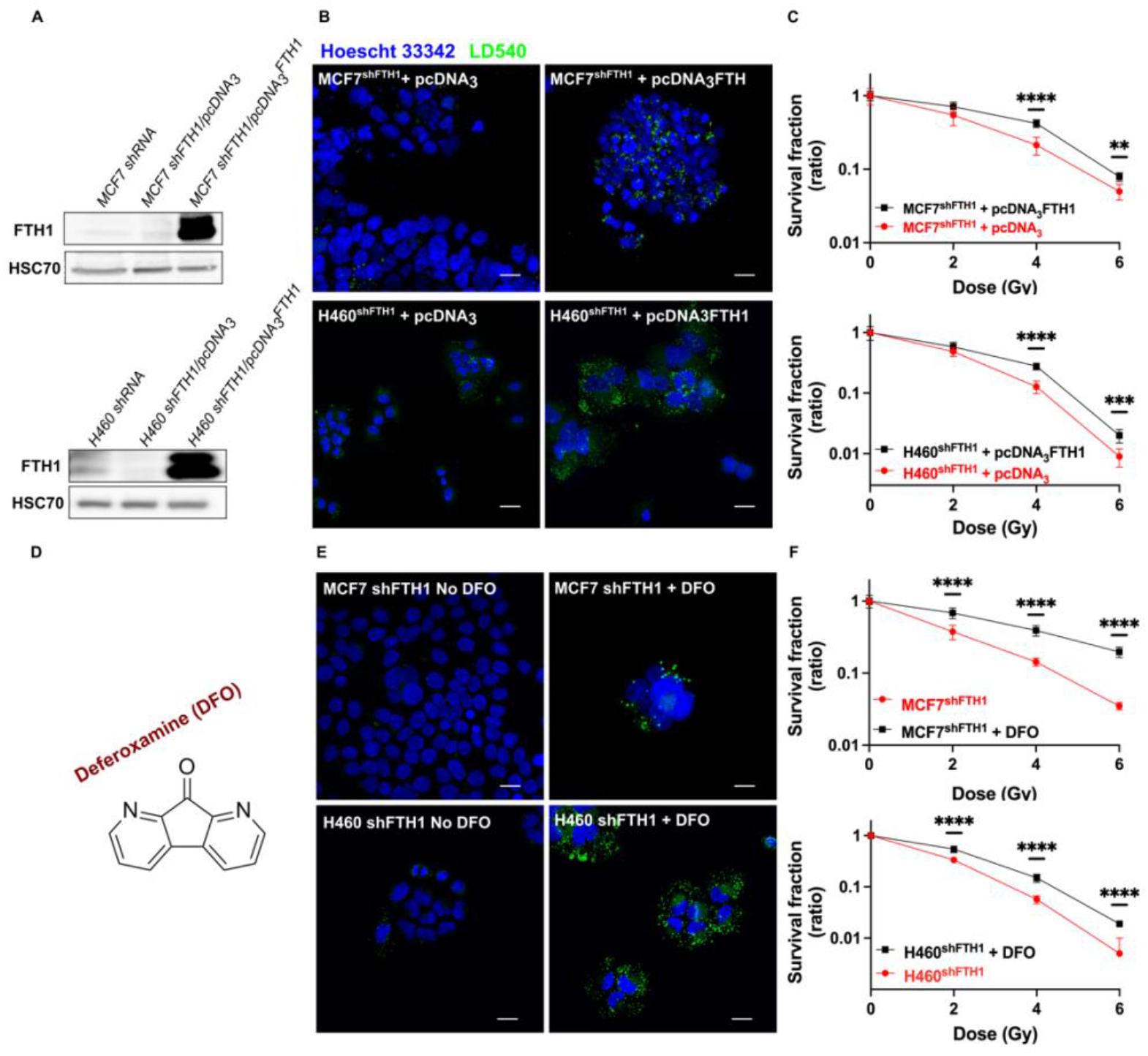
FTH1 reconstitution as well as DFO treatment restore the LD content re-establishing cancer radioresistance. **A**) Western Blotting analysis of FTH1 expression in MCF7 shFTH1/pcDNA_3_FTH1 and H460 shFTH1/pcDNA_3_FTH1. HSC70 was used as loading control. **B)** Z-stack representative confocal fluorescence images of LD detection in MCF7 shFTH1/pcDNA_3_FTH1 and H460 shFTH1/pcDNA_3_FTH1 cells and their H460 shFTH1/pcDNA_3_ and MCF7 shFTH1/pcDNA_3_ controls. (Scale bars 20 μM). (**C**) Survival fractions (in log-linear scale) after FTH1 reconstitution in MCF7- and H460-shFTH1 cells. The iron chelator agent Deferoxamine (DFO), whose chemical structure is reported in **D**, was used for treating MCF7 shFTH1 and H460 shFTH1 for 24 hrs. **(E)** In both cell lines, DFO treatment increased the LD numbers, as showed by confocal microscopy images (Scale bar 20 μM). **(F)** Survival curves (in log-linear scale) of FTH1-silenced MCF7 and H460 cells after DFO treatment. **F**. Error bar represents the means ± SD from three independent experiments. * ≤ 0.05; ** ≤ 0.01; *** ≤ 0.001 and ****≤ 0.0001.

Moreover, such a reconstitution was sufficient to fully restore the LD pool (**Figure 4B**) and to reacquire a higher RR in both cell lines (**Figure 4C**).

To further elucidate the connection between iron and LDs, we used an iron chelator agent, Deferoxamine (DFO), to cope with the iron imbalance due to the FTH1 silencing. DFO is a high-affinity Fe^3+^ chelator and an FDA approved drug used to treat patients with iron overload. H460^shFTH1^ and MCF7^shFTH1^ were treated with DFO for 24 hrs and their LD content was analyzed. LD540 staining on both treated H460^shFTH1^ (H460 shFTH1 + DFO) and MCF7 shFTH1 (MCF7 shFTH1 + DFO) univocally showed that the iron chelation was able to induce LD accumulation and this, in turn, conferred higher survival ability to cells after radiation treatment, as shown by clonogenic assays (**Figure 4F**).

## Discussion

Along with surgery and chemotherapy, radiotherapy represents an important treatment option also in the palliative regimens. Great advance in the understanding of the molecular mechanisms underlying the cancer RR have been done. Nevertheless, this has not translated into a proportional improvement of the therapeutic outcomes because multiple biological factors and their complex interactions capable of negatively affecting the cellular response to ionizing radiation remain to be characterized. Radioresistance exhibited by many cancer cells, especially CSCs, severely limits the effectiveness of the treatments. For this reason, the identification of specific features for targeting RR cells is critical, and it is currently the focus of intense research. Classical CSC markers used to identify the most putative RR cells are still being debated due the high intra- and inter-tumor heterogeneity ^22^ and the cancer cell ability to change during cancer progression and treatments. In recent years, accumulating studies suggested that LDs might be correlated with a CSC phenotype and an elevated tumorigenic potential ^4^. Increased LD amount has been found in various cancer cells with stem-like properties, including colorectal cancer cells ^4^, glioblastoma cells ^7^ and breast cancer cells ^6^.

In the present study, lung (H460), neuroglioma (H4), breast (MCF7), prostate (PC3) and bladder (T24) cancer cells were irradiated with 6 Gy X-rays and left in culture for 3 days in order to select only the RR cells. Surviving cells from all cell lines exhibited an elevated LD content, to which corresponded a cell type-dependent upregulation of the PLIN genes, whose proteins play a role in the formation and structure of LDs. These RR cells also showed differential and cell-specific upregulation of some CSC markers in almost all cell lines. Although we did not screen all the putative stemness markers, our preliminary data indicates that radiation exposure might enrich the heterogeneous population with cells having a stemness-like phenotype, which is in agreement with previous works ^18,20^.

Meanwhile, RR cells with high LD content showed higher ROS levels associated with an increased antioxidant ability, as demonstrated by the genetic upregulation of antioxidant scavenging enzymes, such as SOD and Catalase. However, this behavior was not common to all cell lines, and, in fact, in T24 ROS levels remained unchanged. This suggest that cells from different origin were able to deal with the high dose radiation in different ways, but they all shared the ability to accumulate cytoplasmic LDs. Of note, the presence of cells with high levels of ROS in the not-irradiated cells also displaying high levels of LDs suggest that LDs could serve an antioxidant system being able to buffer the excess of ROS. Indeed, high ROS levels are commonly found in many cancer cells and LDs could contribute to create a tolerable oxidative microenvironment and to better counteract the excess of ROS produced by irradiation.

In support of that, we demonstrated that a higher LD content was a feature already present in the heterogeneous populations, as pre-sorted (LD^high^ and LD^low^) cells displayed differential survival capacity after radiation, with the LD^high^ subpopulation displaying the highest clonogenic response. This indicates that the presence of a higher LD amount was an intrinsic feature of the cells and may represent a selective advantage which might allow resistant cells to survive damages, including oxidative stress induced by ROS production following exposure to ionizing radiation. Many intracellular mechanisms participate in ROS production and the Fenton reaction is one of them. In this reaction, ferrous ion is used as a catalyst to convert H_2_O_2_ into the highly oxidative hydroxyl radical (OH•). Iron is an important player in normal cells because it is involved in many processes and therefore its homeostasis is tightly regulated. However, in cancer cells iron homeostasis is dysregulated and Ferritin, the protein involved in iron storage, has been shown to be elevated in some tumor tissues, thus suggesting that increased iron storage in cancer cells might contribute to cell survival ^11^.

Previous studies showed a crucial role for FTH1 in cancer aggressiveness. In FTH1-silenced MCF7 and H460, cells acquired a mesenchymal phenotype associated with an epithelial to mesenchymal transition and the activation of the CXCR4/CXCL12 signaling pathway ^12^. However, studies on Ferritin and iron roles in radiotherapy are limited. Naz and colleagues demonstrated an upregulation of hepatic ferritin and elevated FTL serum levels in sham-irradiated rats ^10^. In this work, we investigated the effects of X-ray radiation on RR cancer cells in order to determine a possible relationship between FTH1 and LDs. We found that surviving cells in all lines showed an upregulation, although at different extents, of FTH1 protein. Moreover, in both MCF7- and H460-LD^High^ fractions, FTH1 resulted upregulated as compared to the MCF7- and H460-LD^Low^ counterparts. The link between FTH1 and LDs was further confirmed by the reduction of LD accumulation in FTH1-silenced cells. Moreover, FTH1 downregulation associated with the downregulation of transferrin receptor that mediates extracellular iron uptake and the upregulation of ferroportin, responsible for iron release, indicating that most likely iron levels in FTH1-silenced cells were unbalanced. In such conditions, cells were significantly more sensitive to ionizing radiation than the relative controls. These findings show a strong correlation between FTH1 expression and LD content in radioresistant cells and, indirectly, suggest that unbalanced intracellular availability of iron produced effects on lipid pathways, mainly on LD accumulation. These data were then corroborated by restoration of FTH1 expression in silenced H460 and MCF7 cell lines, where we observed a restored LD content together with an increased clonogenic response. Our findings support the idea that the two cell states (LD^High^/FTH1^High^ and LD^Low^/FTH1^Low^) are not irreversible processes, but they are reversible mechanisms where the big player is the cytoplasmic iron pool. Alteration in FTH1 expression induce alterations of intracellular free Fe levels. Excess iron is cytotoxic, mainly because of the production of ROS, and in our study cells with reduced ability to store iron also showed reduced RR. However, in absence of adequate levels of FTH1, treatment with an iron chelator was able to reduce the excess iron inside cells and this caused a significant LD re-accumulation in both FTH1-silenced (MCF7 and H460) cell lines. Once again, re-established LD content and iron storage resulted in increased RR of both cell lines. Therefore, iron homeostasis is strongly correlated to the surviving ability of the RR cells and LDs are important mediators in these processes.

Although the data reported here need to be validated in more physiologically complex systems, they provide novel insights about LD involvement in the radio-resistance of cancer cells and show that this feature is common to different tumor cells analyzed in the present study. Further, our data describe the dynamic interplay between LDs and iron homeostasis, showing that it plays a crucial role in the context of tumor RR. These functional cross-talks need to be more deeply explored in order to determine the potential contribution of other related pathways and organelles in these processes. This would offer the opportunity for a better understanding of the mechanisms behind radiation responses and may suggest novel strategies for incrementing the radiotherapy curative capacity.

Lastly, a common effort has to be put forth in the identification of robust and functional predictive biomarkers to be used to target the most resistant cancer populations by precise treatments, which need to be as specific as possible for the most tumorigenic cells (CSCs/CICs) while preserving, as much as possible, toxicity on the healthy cell population ^19^.

## Materials and Methods

### Cell Cultures and Transfection

MCF-7 human breast adenocarcinoma and H4 neuroglioma cell lines (ATCC) were cultured in DMEM medium (Thermo Fischer Scientific) supplemented with Fetal Bovine Serum (FBS) 10% (Thermo Fischer Scientific), Pen/Strep 1% (Thermo Fischer Scientific). H460 human non-small lung cancer cells (ATCC) were cultured in RPMI 1640 (Thermo Fischer Scientific) medium supplemented with 10% FBS and 1% Penicillin-Streptomycin (Thermo Fischer Scientific). T24 bladder carcinoma cell line (ATCC) was cultured in McCoy’s medium (Thermo Fischer Scientific) supplemented with FBS 10% (Thermo Fischer Scientific), Pen/Strep 1% (Thermo Fischer Scientific) and Hepes 1% (Thermo Fischer Scientific). PC3 prostate adenocarcinoma cells (ATCC) were cultured in F-12K medium (Thermo Fischer Scientific), supplemented with FBS 10% (Thermo Fischer Scientific), Pen/Strep 1% (Thermo Fischer Scientific). All these cell lines were maintained at 37°C in a humidified 5% CO_2_ atmosphere and cultured following ATCC recommendations.

Lentiviral transduced MCF7 and H460 were kindly provided by the laboratory headed by Prof. Francesco Saverio Costanzo at the University Magna Graecia of Catanzaro. Both cell lines were stably transduced with a lentiviral DNA containing either an shRNA that targets the 196–210 region of the FTH1 mRNA (sh29432) (MCF-7shFTH1, H460shFTH1) or a control shRNA without significant homology to known human mRNAs (MCF-7shRNA, H460shRNA). MCF-7shRNA and MCF-7shFTH1 were cultured in DMEM medium (Thermo Fischer Scientific) supplemented with FBS 10% (Thermo Fischer Scientific), Pen/Strep 1% (Thermo Fischer Scientific), puromycin 1μg/ml (Sigma-Aldrich). H460shRNA and H460shFTH1 were cultured in RPMI 1640 (Thermo Fischer Scientific) medium supplemented with 10% FBS and 1% Penicillin-Streptomycin (Thermo Fischer Scientific), puromycin 1μg/ml; All cell lines were maintained at 37°C in a humidified 5% CO_2_ atmosphere.

### Radiation Treatment and Clonogenic Assay

Irradiation has been carried out using a Multi Rad 225kV irradiator. Cells, seeded at a density of 3.5 × 10^5^ and 1.0 × 10^6^ cells for 0 and 6 Gy respectively, were irradiated with 6 Gy at room temperature and left in culture for 72 hours in order to get only surviving cells at the end of the culturing time. Fresh medium was replaced every day.

Cell survival was evaluated using a standard colony forming assay. H4 LD^High^ and LD^Low^, H460 LD^High^ and LD^Low^, MCF7 LD^High^ and LD^Low^, PC3 LD^High^ and LD^Low^, T24 LD^High^ and LD^Low^, H460 shRNA, H460 shFTH1, H460 shFTH1 + DFO, H460 shFTH1/pcDNA_3_, H460 shFTH1/pcDNA_3_FTH1, MCF7 shRNA, MCF7 shFTH1, MCF7 shFTH1 + DFO, MCF7 shFTH1/pcDNA_3_ and MCF7 shFTH1/pcDNA_3_FTH1 populations were collected soon after sorting. Cells were seeded into six well plates (Corning) at a density of 2 × 10^2^ −1 × 10^4^ cells/well, irradiated (2, 4 and 6 Gy single dose) with a Multi Rad 225kV irradiator and incubated for 7-12 days at 37°C in a humidified atmosphere with 5% CO_2_. Following incubation, colonies were fixed in 100% ethanol and stained using a 0.05% crystal violet solution. Only the colonies with more than 35 cells were counted. Surviving fractions were calculated after correction for plating efficiency of control cells. At least three independent experiments, each in duplicate, have been performed for the above-mentioned cell samples.

### Cell Sorting

T24, MCF-7, H460, H4, PC3 cell suspensions were washed in Phosphate-Buffered Saline (PBS) (Thermo Fischer Scientific). Cells were then stained with LD540 for 10 min at 37°C in the incubator. The excess of dye was washed away with PBS and cells were resuspended in sorting buffer (PBS Ca/Mg-free, BSA 0,5%, EDTA 2 mM and Hepes 15mM).

Cells were sorted in two populations (LD^High^ and LD^Low^) using a FACSAria Fusion (BD Bioscience). Sorting gates were established based on the 10% most bright and 10% most dim subpopulation. All cell sorting experiments have been carried out within 1 hour upon sorting to avoid that sorted cells could start becoming heterogeneous again.

### Lipid Droplet Staining

Depending on the project needs, LD content was assessed by staining cells with two different dyes: LD540 and Nile Red. For FACS measurements, 4 × 10^5^ cells have been harvested, washed with PBS and then stained with 0.1 μg/ml LD540 or 1/500 (from a saturated stock solution in acetone) Nile Red. Stained cells were analyzed at the FACS Canto II (BD Bioscience). Instead, for the confocal microscopy analysis, 4 × 10^3^ cells have been cultured on a 35mm Glass Bottom Dishes (MatTek Life Science) and then, fixed with PFA 4%. After washing out the PFA, fixed cells were stained with 0.1 μg/ml LD540 and 1 μg/ml Hoechst 33342 (Thermo Fischer Scientific). Cells were imaged by a Leica SP5 or a Zeiss LSM710 confocal microscope systems.

### ROS Staining

Intracellular ROS content was measured by freshly prepared chloromethyl dichlorodihydrofluorescein diacetate (CM-H_2_DCFDA, Thermo Fisher Scientific) dye resuspended in anhydrous dimethyl sulfoxide (Thermo Fisher Scientific). Briefly, 4 × 10^5^ cells were collected and washed three times with PBS Ca^+^/Mg^+^-free 1X and soon after incubated with 3.5 μM CM-H_2_DCFDA in pre-warmed Hank’s balanced salt solution (HBSS, Thermo Fischer Scientific) for 20 min at 37°C, in the dark. The samples were analyzed, after having washed them with PBS, by using a FACSCanto II flow cytometer (BD Biosciences).

### Lipid Droplet and ROS co-staining

4 × 10^5^ cells were harvested, washed with PBS 1X and soon after stained with 1/500 (from a saturated stock solution in acetone) of Nile Red for 20 min at 37°C in the dark. Stained cells were washed three times and then incubated with 3,5 μM of CM-H_2_DFCDA in HBSS for 20 min at 37°C in the dark. After one wash in PBS 1X, cells were analyzed using a FACS Canto II (BD Bioscience).

### Antibodies and Western Blot Analysis

H4 0 and 6 Gy, H460 0 and 6 Gy, MCF7 0 and 6 Gy, PC3 0 and 6 Gy, T24 0 and 6 Gy, MCF7 shRNA, MCF7 shFTH1/pcDNA_3_, MCF7 shFTH1/pcDNA_3_FTH1, H460 shRNA, H460 shFTH1/pcDNA_3_ and H460 shFTH1/pcDNA_3_FTH1 cells were washed twice with cold PBS and incubated for 20 min with 300 μL of 1X Ripa Buffer (Cell Signaling) additioned with HaltTM Protease Inhibitor Single-Use Cocktail, (Thermo Fisher Scientific) and HaltTM Phosphatase Inhibitor Single-Use Cocktail (Thermo Fisher Scientific), both diluted 1:100. Cells were then transferred to tubes and, after centrifugation at 14000xg at 4°C for 20 minutes, the supernatants were collected. Protein concentration was measured by BCA Protein assay kit (Thermo Fisher Scientific) at 562 nm using BSA to produce a standard curve. For protein analysis, 15 μg of whole cell extracts for each sample were electrophoresed under reducing condition in 10% SDS-polyacrylamide gels and then electrophoretically transferred onto PVDF membrane filters (Bio-Rad Laboratories), using Trans-Blot Turbo Transfer System (Bio-Rad Laboratories, Hercules, CA, USA). In order to prevent the non-specific antibody binding, blots were blocked for 1 hr with BSA blocking buffer, 5% in PBS, with 0,1% Tween-20 (TWEEN 20 Bio-Rad Laboratories). Membranes were washed with PBS-0.1% Tween and incubated with antibodies in blocking solution overnight at 4 °C. Antibody used was a rabbit anti H-ferritin (1:200; Santa Cruz Biotechnology, Texas, USA). PBS-0.1%Tween-20 was used to remove the excess of primary antibody and then the membranes were incubated in blocking solution with goat anti-mouse IgG-HRP (1:2000, Santa Cruz Biotechnology) secondary antibody. Subsequently, blots were rinsed with 0.1% PBS-Tween and developed with Clarity Western ECL Substrate (Bio-Rad Laboratories) using Amersham Imager 680. Protein levels were analyzed by ImageJ 1.52p software.

### RNA isolation and Real-Time PCR (qRT-PCR)

Total RNA was isolated from 6 Gy irradiated and non-irradiated cells, LD^High^ and LD^Low^ sorted cells, MCF-7 shRNA and MCF-7 shFTH1 as well as H460 shRNA and H460 shFTH1 using the High Pure RNA isolation kit (Roche) according to the manufacturer’
ss instructions. All the RNA samples were treated with DNase-1 to remove any contaminating genomic DNA and the RNA purity was checked spectroscopically. Then, 1 μg of purified RNA was reverse transcribed using RT^2^ First Strand Kit (Qiagen) according to the manufacturer’s instructions.

Gene expression analysis was assessed by Real-Time PCR (qRT-PCR) using the cDNA obtained from the cell samples above reported.

20 ng of cDNA was amplified in 15 μl of reaction mix containing Power SYBR Green PCR Master mix (ThermoFisher Scientific), 20 pmol of each primer pair and nuclease-free water on a StepOne Plus System (ThermoFisher scientific). The thermal profile consisted of 1 cycle at 95 °C for 10 min followed by 40 cycles at 95°C for 15 sec, 60°C for 1 min. Relative gene expression was normalized to that of the gene encoding the human GAPDH which served as an internal control. Data analysis was performed using the 2-ΔΔCt method.

### Widefield and Confocal Microscopy

T24, H4, PC3, MCF7, MCF7 shRNA, MCF7 shFTH1, H460, H460 shRNA and H460 shFTH1 were seeded and stained with LD540 as reported in the Lipid Droplet Staining section. Zeiss LSM710 and Leica SP5 microscopes, both equipped with a 40X and 63X, were used to image LDs.

### Image Analysis

Z-stack images of LD540 stained cells were taken using a Leica SP5 confocal-laser-scanning microscope equipped with a 40x oil immersion i-Plan Apochromat (numerical aperture 1.40) objective. LD540 were visualized using the 488 nm line of an Argon laser and a 505-530 nm BP filter. 12-bit images were acquired and post processed for the LD quantification. Briefly, the background was subtracted using ImageJ’s Rolling ball radius tool. The images were further processed with Gaussian filter, thresholded and segmented with Find Maxima tool. Finally, images were analyzed with Analyze Particles tools. All the image processing was performed automatically with constant settings using in house developed macro for Fiji generously provided by Dr. Damir Krunic.

Student’s t-test with unequal variances was used for the calculation of statistical significances. Differences of two groups with P values below 0.05 were considered statistically significant.

### FTH1 Reconstitution

MCF7 shFTH1 and H460 shFTH1 cells were seeded in six-well plates at 3 × 10^5^ cells/well and grown overnight prior to transfection.

All plasmids were transfected with Lipofectamine 3000 transfection reagent (Thermo Fisher Scientific) following manufacturer’s instructions. FTH1 reconstitution was performed using 2,5 μg/μl of the expression vector containing the full length of human FTH1 cDNA (pcDNA3/FTH1) (MCF-7 shFTH1/pcDNA_3_FTH1 and H460 shFTH1/pcDNA_3_FTH1) while 2,5 μg/μl of pcDNA_3_ plasmid was used as negative control (MCF-7 shFTH1/pcDNA_3_ and H460 shFTH1/pcDNA_3_). Transfection efficiency was tested by western blot and qPCR after 48 hrs. All transfection experiments were repeated in triplicate.

### Deferoxamine Treatment

MCF-7-Wt, MCF-7-shRNA, MCF-7-shFTH1, H460-Wt, H460-shRNA and H460-shFTH1 cells were seeded in 100 mm^2^ petri dishes (Corning) at a concentration of 1,5 × 10^6^ cells/plate containing 10 mL of DMEM or RPMI-1640 (supplemented with 10% FBS) and incubated for 24 hrs. Then, cells were treated with 50 μM DFO (Deferoxaminemesylate salt). Cells cultured in normal medium were used as control. After 24 hrs of treatment, cells were collected and used for ROS and LD detection.

### Statistics

All data here presented are shown as mean values ± SD of the irradiated or “treated” samples relative to the untreated control. Statistical and data analysis was carried out using GraphPad Prism 9 software. Statistical differences between treated and untreated samples were assessed by T-Test and one-way ANOVA. The threshold for statistical significance was set to P = 0.05.

## Acknowledgments

We gratefully acknowledge the Imaging and FACS facilities of the DKFZ for their precious support. LT has received funding from AIRC and from the European Union’s Horizon 2020 Research and Innovation Programme under the Marie Skłodowska-Curie grant agreement No 800924.

## Author Contributions

L.T. and J.S. designed and coordinated the whole project; L.T., M.G.M., F.P. and J.S. designed the experiments; L.T., M.G.M., J.J., I.A., R.H., and C.N. performed and analyzed all clonogenic assays M.G.M., I.A. and F.S.C. performed and supervisioned the FTH1 silencing as well as the FTH1 reconstitution; L.T., F.P., J.F.G. and D.G.C. carried out the confocal analyses; M.G.M. and I.A. performed the DFO treatment; L.T., M.G.M., F.P., D.G.C. and J.J. performed the data analysis; L.T., M.G.M and F.P. wrote the manuscript; L.T., F.P., F.S.C. and J.S. revised the manuscript.

## Supplementary Information for

### This PDF file includes

Figure S1 to S4

Table S1

**Figure S1:**
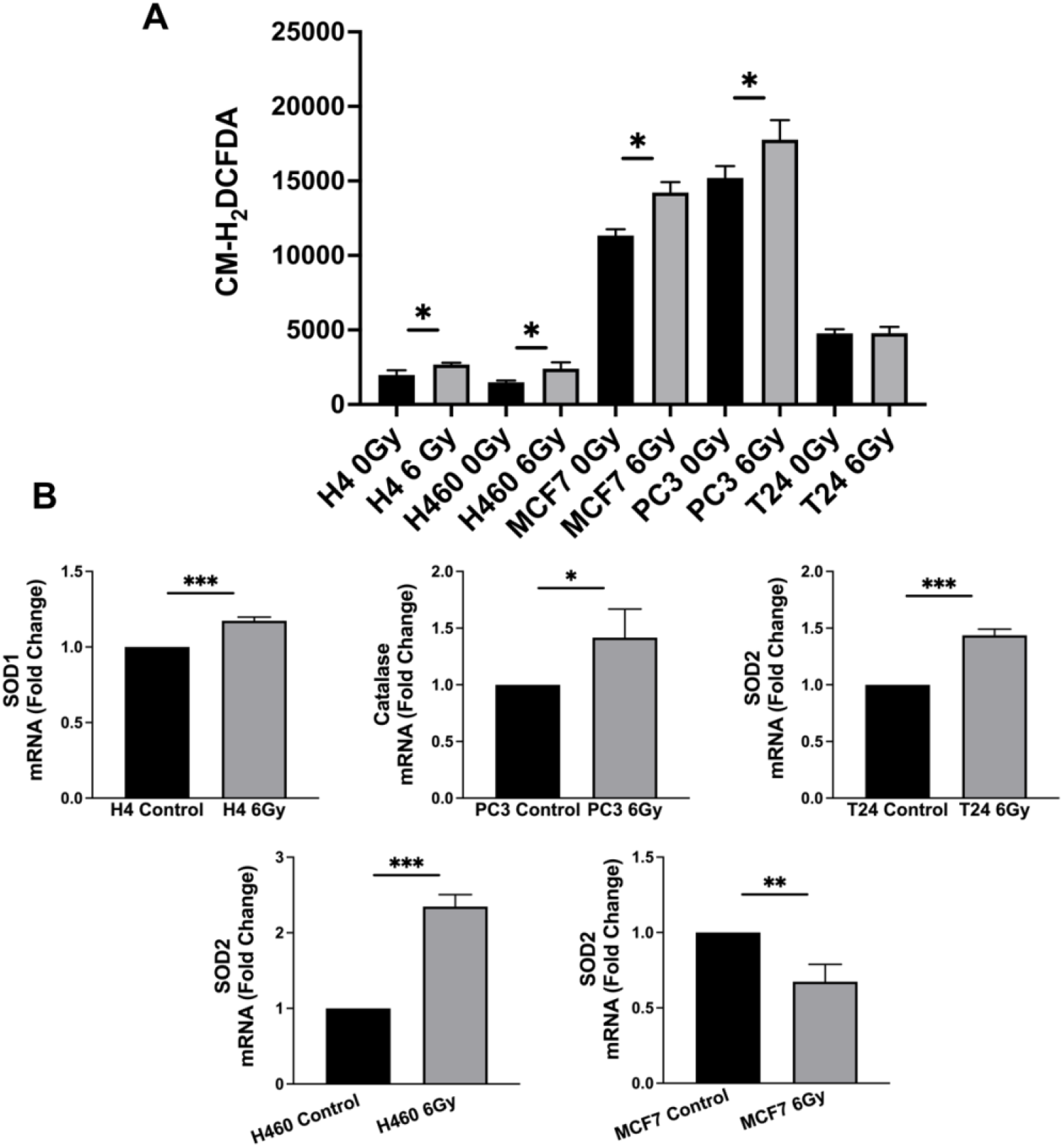
ROS evaluation in 6 Gy radioresistant cancer cells. **A)** ROS staining by CM-H_*2*_DCFDA probe *was* performed in 6 Gy tr*e*ated and untreated H4, H460, MCF7, PC3 and T24 cancer cell lines. **B)** Gene expression analysis of genes related to oxidative express responses by RT-qPCR. SOD1, SOD2, Catalase and GPX have been evaluated and only the significative genes are here reported. All data represent the means ± SD from three independent experiments. * ≤ 0.05; ** ≤ 0.01; *** ≤ 0.001 and ****≤ 0.0001.

**Figure S2:**
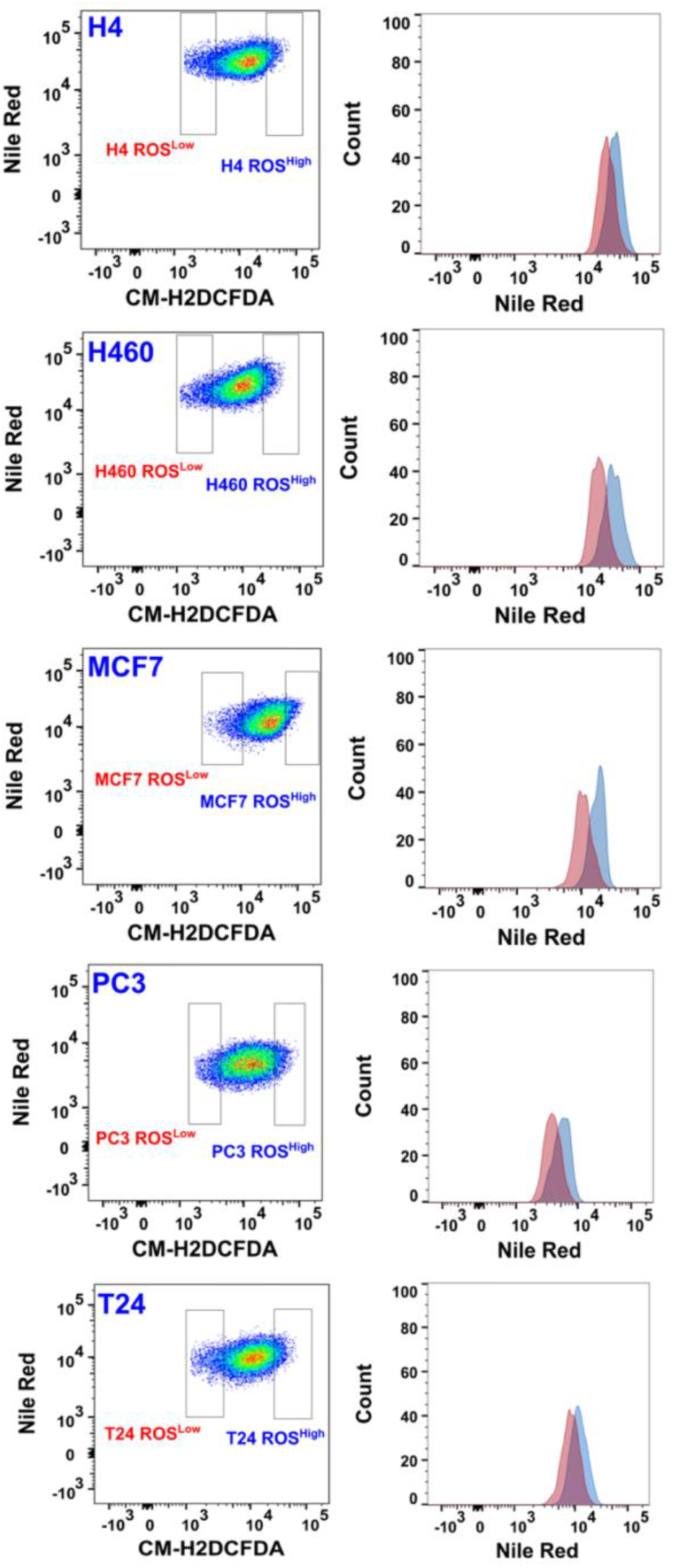
ROS and Lipid Droplet Double Staining. Heterogeneous H4, H460, MCF7, PC3 and T24 cancer were double stained with CM-H_2_DFCDA and Nile Red for ROS and Lipid Droplet evaluation, respectively. In the right panel, a representative FACS plot of cells showing that the highest amount of ROS also corresponded to the highest number of LDs.

**Figure S3:**
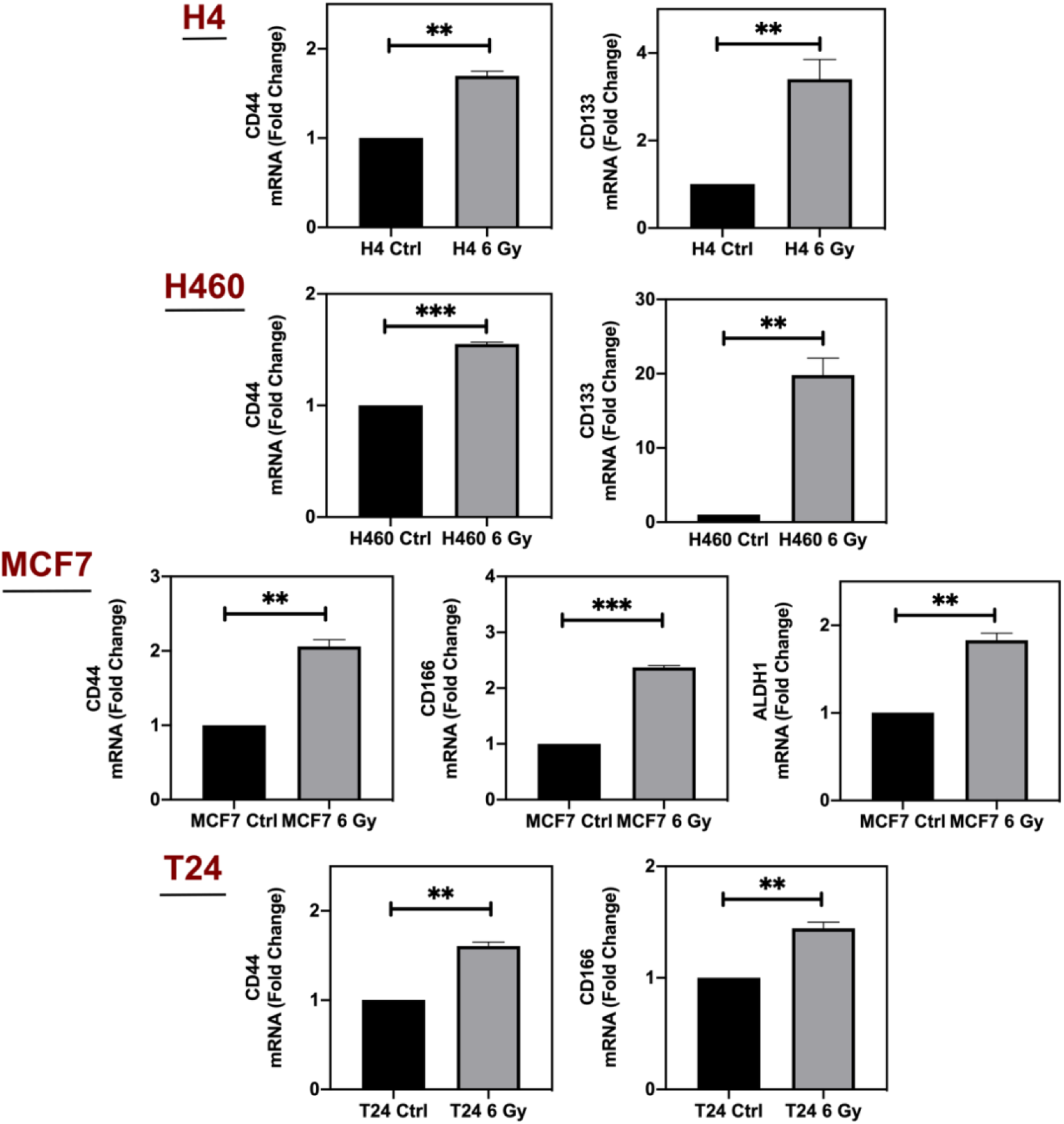
CSC Marker Evaluation in 6 Gy Radioresistant Cells. CD44, CD133, CD166 and ALDH1 mRNA expression in 6 Gy radioresistant H4, H460, MCF7, PC3 and T24 cancer cells as assesed by RT-qPCR. Only the significative genes are here reported. All data represent the means ± SD from three independent experiments. * ≤ 0.05; ** ≤ 0.01; *** ≤ 0.001 and ****≤ 0.0001.

**Figure S2:**
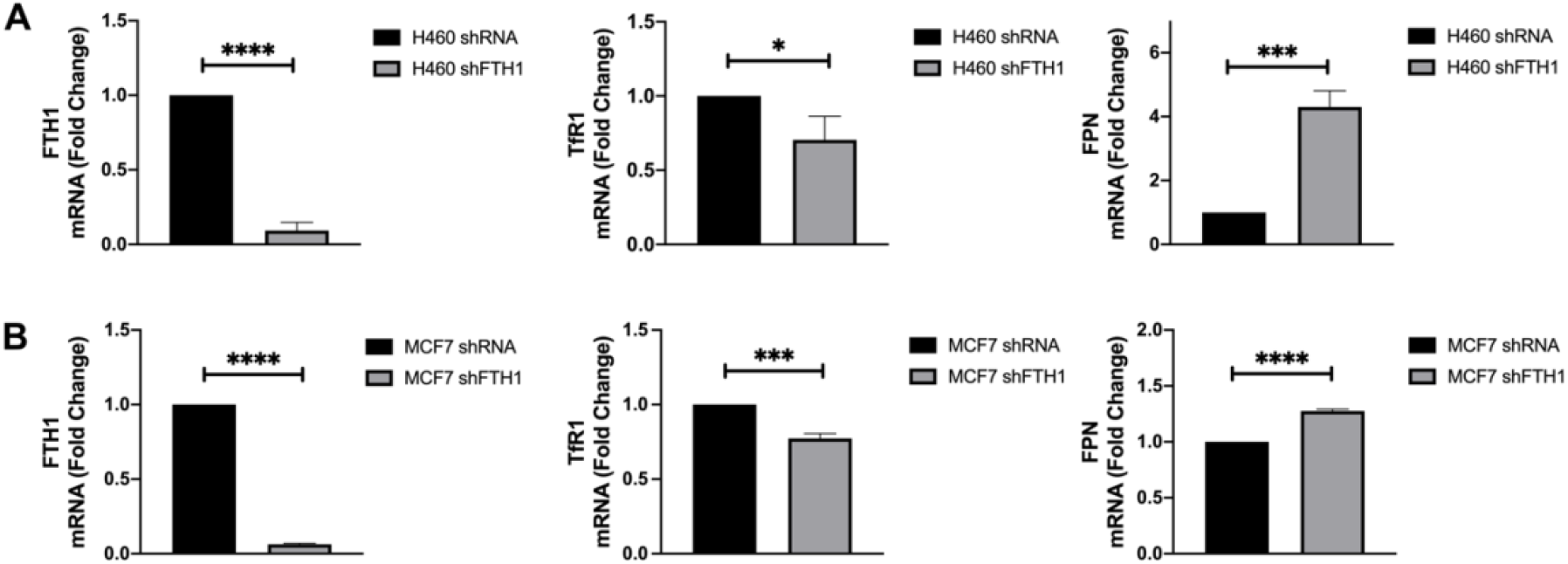
FTH1 silencing in H460 and MCF7 causes TfR1 downregulation and FPN upregulation. RT-qPCR analysis of H460 and MCF7 silenced for FTH1 shows TfR1 mRNA downregulation and FPN upregulation in both cell systems. All data represent the means ± SD from three independent experiments. * ≤ 0.05; ** ≤ 0.01; *** ≤ 0.001 and ****≤ 0.0001.

**Table S1.**
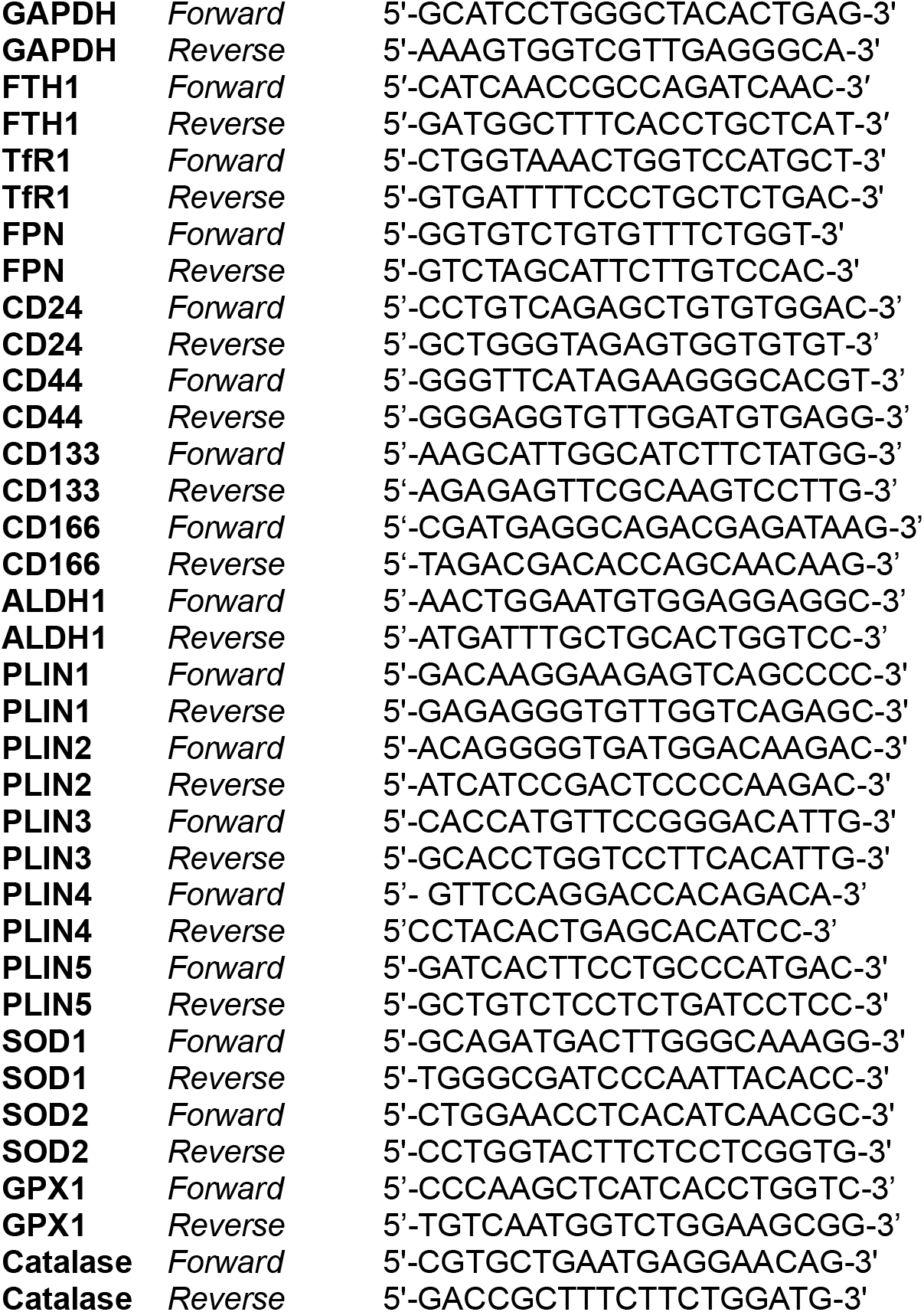
Sequence of qRT-PCR primers used in this study.

